# Freezing displayed by others is a learned cue of danger resulting from co-experiencing own-freezing and shock

**DOI:** 10.1101/800714

**Authors:** Andreia Cruz, Mirjam Heinemans, Cristina Marquez, Marta A. Moita

**Author notes:** Senior author (Marta A. Moita).

## Abstract

Social cues of threat are widely reported [1–3], whether actively produced to trigger responses in others, such as the emission of alarm calls, or by-products of an encounter with a predator, like the defensive behaviors themselves, such as an escape flight [4–14]. Although the recognition of social alarm cues is often innate [15–17], in some instances it requires experience to trigger defensive responses [4,7]. One mechanism proposed for how learning from self-experience contributes to social behavior is that of auto-conditioning, whereby subjects learn to associate their own behaviors with the relevant trigger events. Through this process the same behaviors, now displayed by others, gain meaning. [18,19 but see: 20]. Although it has been shown that only animals with prior experience with shock display observational freezing [21–25] suggesting that auto-conditioning could mediate this process, evidence for this hypothesis was lacking. Previously we found that when a rat freezes, the silence that results from immobility triggers observational freezing in its cage-mate, provided the cage-mate had experienced shocks before [24]. Hence, in our study auto-conditioning would correspond to rats learning to associate shock with their own response to it – freezing. Using a combination of behavioral and optogenetic manipulations, here we show that freezing becomes an alarm cue by a direct association with shock. Our work shows that auto-conditioning can indeed modulate social interactions, expanding the repertoire of cues that mediate social information exchange, providing a framework to study how the neural circuits involved in the self-experience of defensive behaviors overlap with the ones involved in socially triggered defensive behaviors.

## RESULTS

Various studies using different paradigms have shown that prior experience with shock is required for a robust display of observational freezing, [21–23,25]. In a previous study we examined observational freezing by placing pairs of cage-mate rats in a two-chambered box, separated by a partition that allowed rats to interact. One of the rats in the dyad, the conditioned demonstrator, froze upon a tone previously paired with footshock. The other rat in the dyad, the observer cage-mate that had never been exposed to the tone, responded to the freezing of the demonstrator by freezing too, but only if it had previously experienced unsignaled shocks [24]. In the present study we set out to investigate how prior experience with shock facilitates observational freezing. We first hypothesized that the stress of receiving unsignaled footshocks could sensitize neural circuits that regulate defensive responses, causing observer animals to respond with increased intensity to otherwise neutral or novel stimuli [26]. To test this hypothesis we subjected observer rats to a different type of uncontrollable emotional stressor, the forced swim session (FS) [27] and compared the level of observational freezing of these rats to those of observer rats subjected to our standard conditioning session: three unsignaled shocks about three minutes apart (from now on “spaced shocks”, SS) (Figure 1A-B). We first verified the stress response induced by our forced swim and spaced shock sessions, by measuring circulating levels of the stress response hormone corticosterone [28,29]. Despite the high levels of corticosterone triggered by the FS session (Figure S1A-B), FS observer rats did not respond to the freezing of demonstrators. In contrast, as previously shown [24], SS observer rats displayed robust observational freezing, even if showing significantly lower levels of corticosterone. In this experiment demonstrators paired with FS-observers and demonstrators paired with SS-observers showed indistinguishable levels of freezing (Figure S1C). Still, due to the nature of the social interaction where individuals influence each other, with the behavior of the demonstrator affecting the observer and vice versa, to directly compare the response of observer rats across conditions, we normalized the freezing of the observer by the freezing of the demonstrator ((Freezing Demonstrator – Freezing Observer)/(Freezing Demonstrator + Freezing Observer)). A normalized freezing score of 1 reflects freezing only by the demonstrator; a score of 0 reflects both rats showing similar level of freezing; and −1 corresponds to freezing only by the observer. This ensures that any difference in demonstrators’ behavior across conditions is accounted for. Figure 1C shows that SS observer rats show normalized observational freezing close to zero (median: 0.01889± 1.129) whereas FS rats close to one (median: 0.8801± 0.6693). A Mann-Whitney U test revealed a significant difference between groups (U= 6, p<0.0001). These results fail to support the hypothesis that stress induced sensitization by itself underlies observational freezing, in line with a prior study using mice [21].

**Figure 1.**
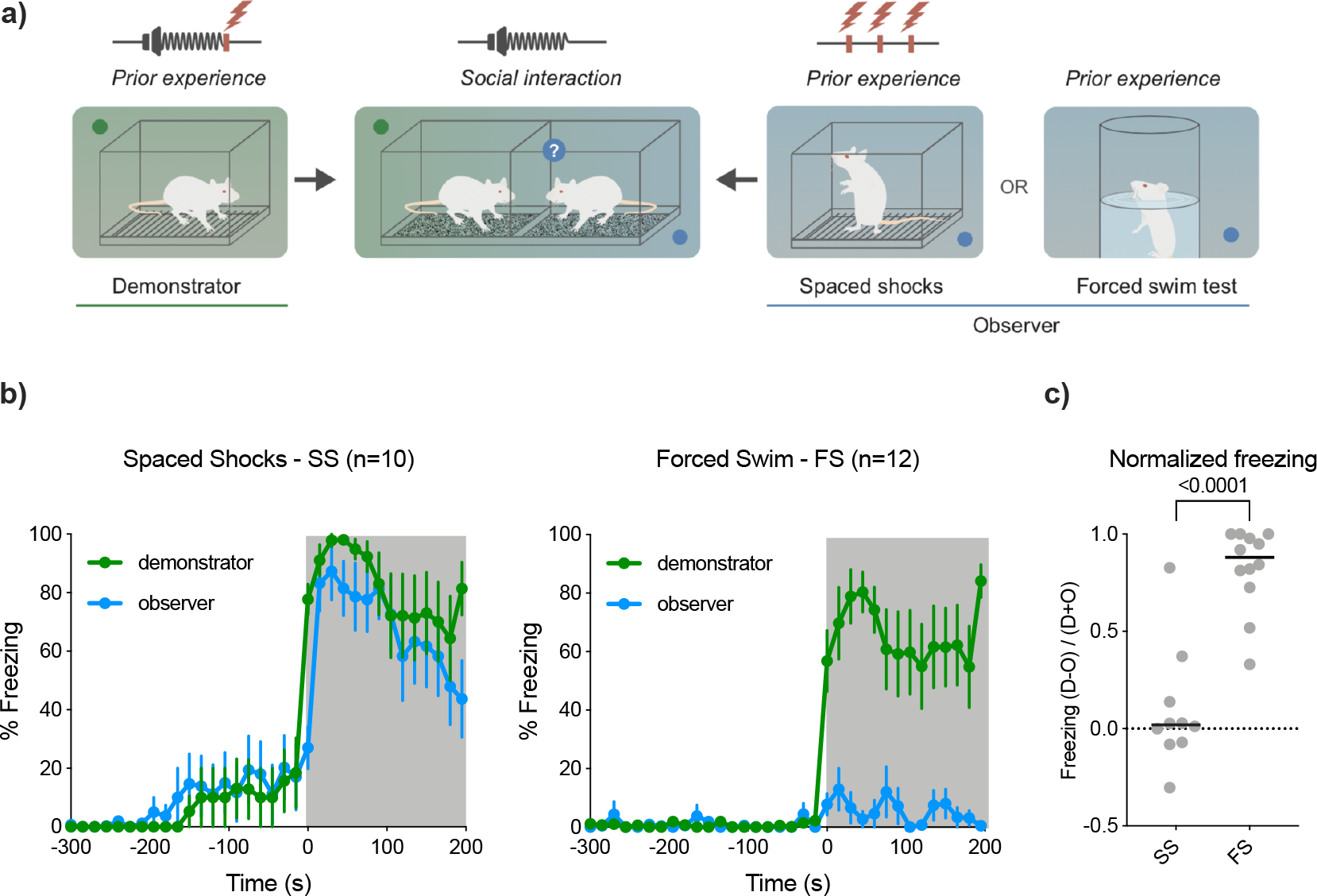
*Stress alone does not drive observational freezing* **a)** Schematic of the behavioral paradigm used to study observational freezing. **b)** Proportion of time spent freezing over time, by pairs of demonstrators and observers during the social interaction. Shaded area corresponds to time after tone presentation. Left panel corresponds to pairs of animals with observers that received SS (n= 10). Right panel corresponds to pairs of animals with observers that experienced FS (n=12). Mean ± S.E.M. **c)** Freezing values per pair, during social interaction, normalized by the freezing of the demonstrator. Line denotes median values. ****p< 0.0001.

### Experiencing freezing is necessary for robust observational freezing

It has been hypothesized that experiencing shocks modulates observational freezing through some form of conditioning [22,25], stress induced sensitization [22] or because animals can recognize similar experiences in others [21]. When exposed to spaced shocks, rats experience pain [30], learn the association between the context in which shocks were delivered and the aversive stimulus [31], and experience their own freezing [32], which could potentially become associated with shock, a form of auto-conditioning [18,19,22,25]. To unravel which of these experiential components is important for the display of observational freezing, we subjected observer rats to different shock protocols that allowed testing these components incrementally, and subsequently compared their freezing in the social interaction session (summarized in Figure 2A and Methods). Rats that received immediate shocks (IS), a paradigm known as the Immediate Shock Deficit, where rats are placed in a novel chamber, immediately shocked and removed [31], did not experience freezing nor learned about the threat [31], but still exhibited the typical unconditioned responses of jumping and squeaking [32], reflecting the aversive nature of the experience. In the delayed shocks (DS) protocol, rats did not experience freezing at the time of the shock, but learned the association between context and aversive shocks as measured by their learned freezing to a later exposure to the context in which they received shock [31] (Figure S2A). Finally, the third group received spaced shocks (SS) as before, and thus experienced a painful stimulus, contextual threat learning and their own freezing response. Importantly, there were no significant differences in the corticosterone levels of animals exposed to these protocols (Figure S2B). We found that only observer animals in the SS group, i.e. that experienced freezing during exposure to shock, displayed robust levels of observational freezing (Figure 2B). The lack of freezing by the observers that experienced immediate shocks has a buffering effect [33] on the behavior of their demonstrators, dampening their response to the threatening tone (Figure S2C). Thus, as before we normalized the observer’s freezing by the freezing of the demonstrator, and performed a Kruskal-Wallis test on the normalized freezing score. We found differences between groups (H= 20.53, p< 0.0001) with post-hoc analysis revealing that IS group was different from SS (p<0.0001) and from DS (p= 0.0046) (Figure 2C). The normalized freezing in the SS and DS groups was not statistically different (p= 0.4701). Closer inspection of normalized freezing scores of the DS group revealed a widespread distribution, with some rat pairs showing high scores and others showing low scores. Hence, we performed a median split of the normalized freezing scores that divided our population in two groups: one where the freezing ratio is close to zero (freezers), in which both observer and demonstrator animals were freezing; and another where the demonstrator froze while the observer did not (non freezers). Figure 2D depicts the mean traces for the time spent freezing during the social interaction, for DS dyads, split into freezers and non-freezers. The comparison of normalized freezing scores separating these two groups shows that DS-freezers did not differ from SS-rats, whereas non-freezers did (Figure S2D). When examining the levels of freezing during baseline, we found that observer rats of the freezers group froze more than those of the non-freezers group (Figure S2E). In addition, in 6/8 dyads the freezer observers started freezing first and/or froze more than demonstrators during baseline, suggesting that freezing in the dyad, during baseline is triggered by the observer rat. This may be explained by generalization of threat learning between conditioning and test chambers [31], since unsignaled shocks (experienced by observers) result in contextual conditioning [31,34], a condition more prone to generalization [35–38] than the signaled shocks experienced by demonstrators [38,39].

**Figure 2.**
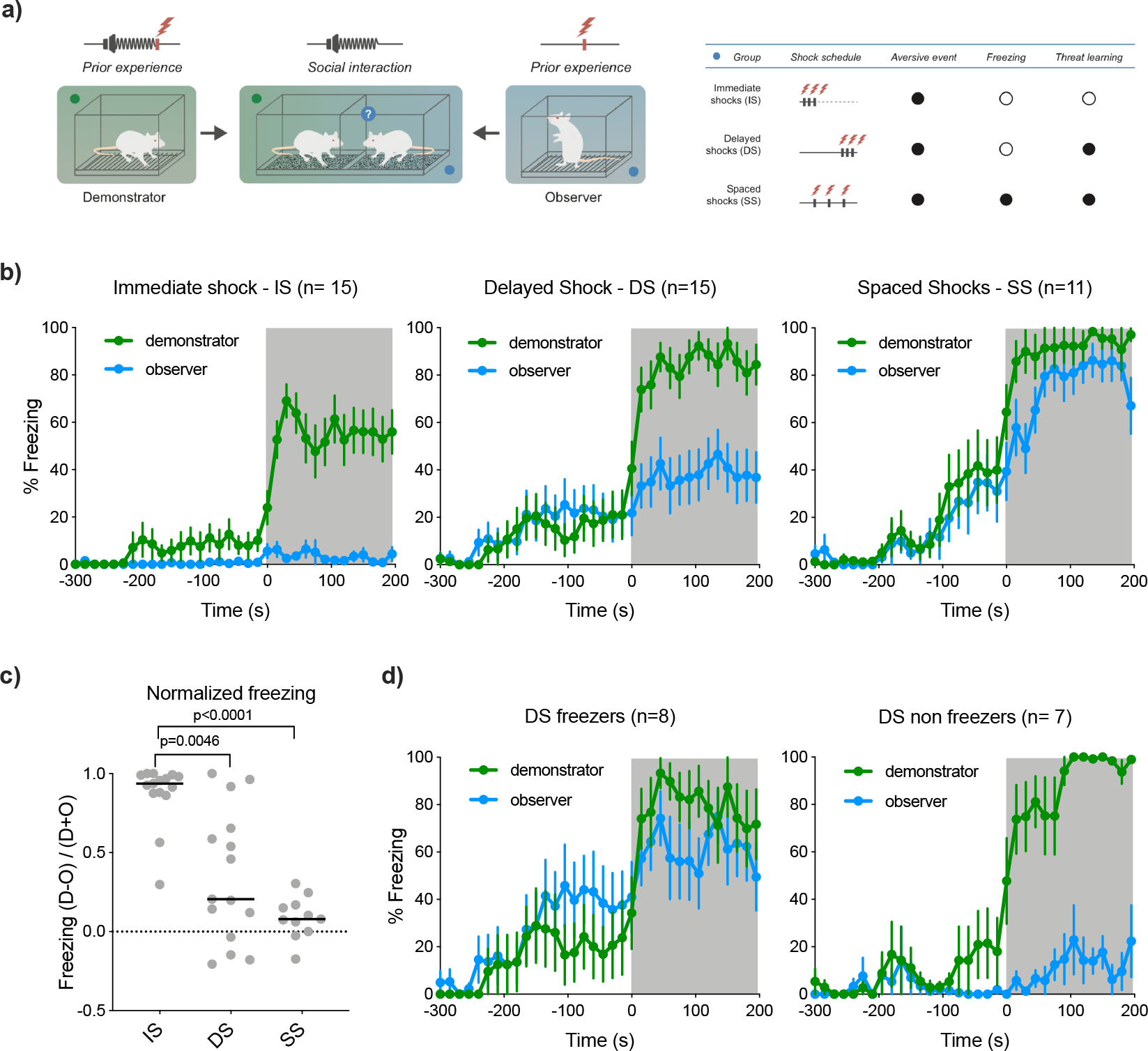
*Manipulation of the schedule of shock delivery to observers* **a)** Left: schematic of the behavioral protocol for the manipulation of experience with shock. Right: Details about the experience of each group of observers. **b)** Proportion of time spent freezing over time, by pairs of demonstrators and observers during the social interaction. Shaded area corresponds to time after tone presentation. Top-left panel corresponds to pairs of animals with observers that received SS (n= 11). Top-right panel corresponds to pairs of animals with observers that experienced IS (n=15). Bottom-left panel corresponds to pairs of animals with observers that experienced DS (n=15). Mean ± S.E.M. **c)** Freezing values per pair, during social interaction, normalized by the freezing of the demonstrator. Line denotes median values. **p<0.01; ****p<0.000.1. **d)** Proportion of time spent freezing over time, by pairs of demonstrators and observers that experienced DS, during the social interaction. Observers were split into freezers (Left: n=8) and non-freezers (right: n=7) by the median of the normalized freezing per pair. Shaded area corresponds to time after tone presentation. Mean ± S.E.M.

Taken together these results show that solely experiencing painful shocks does not facilitate observational learning. However, threat learning can result in generalization, defined by the display of defensive responses in a neutral context after being conditioned [38], contributing in part to the freezing of observers during social interactions. Still observational freezing could be distinguished from contextual generalization as observers that did not generalize, i.e. were not freezing before the demonstrator or during baseline, did not respond with freezing once the demonstrators started displaying this response. Once again, only animals that received spaced-shock (SS) and experienced freezing displayed robust observational freezing.

### Experiencing freezing is not sufficient to drive observational freezing

The previous experiments show that experiencing freezing, triggered by shock, is required for the display of robust observational freezing. Next, we investigated whether experiencing freezing in the absence of painful stimuli or threat learning, could lead to robust observational freezing. To this end, we used two different stimuli that can induce innate freezing, but are not painful. We exposed observer rats to 2MT (an odor derived from TMT a component of fox feces that does not induce threat learning [40]), or to a visual looming stimulus (expanding black circle) whose ability to drive threat learning remains untested [41–44] (Figure 3A). Exposure to looming or 2MT took place in enclosed arenas ensuring that the appropriate response was freezing, as there was no possibility to escape or hide [41,44] (Figure 3B). While 2MT induced weaker freezing responses and no contextual threat learning, exposure to looming stimuli induced robust freezing and low threat learning, whereas spaced shocks induced both robust freezing and threat learning (Figure S3). Only animals that received SS showed robust observational freezing during the social interaction (Figure 3C). A Kruskal-Wallis test was used to compare the normalized scores of freezing between treatments (H= 19.39; p<0.0001). Post-hoc tests revealed that the SS group is different from both the Looming group (p= 0.008) and 2MT group (p<0.0001), which did not differ from each other (p=0.2762) (Figure 3D).

**Figure 3.**
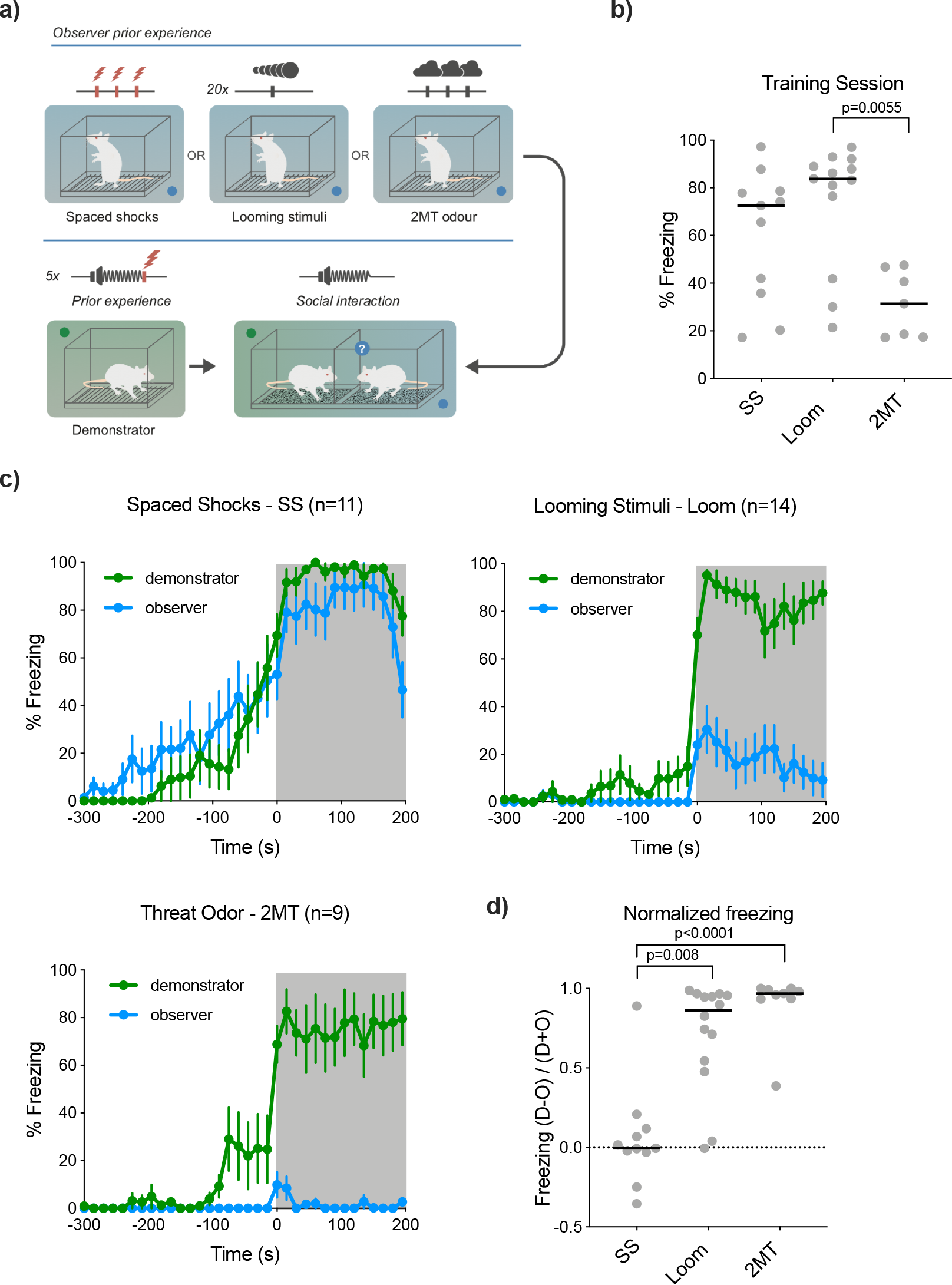
*Manipulation of the experience of freezing* **a)** Schematic of the behavioral protocol for the manipulation of experience with freezing. **b)** Comparison of total time spent freezing, by observers, during the exposure to freezing inducing stimuli. SS (n=11), Loom (n= 13) and 2MT (n=7). A Kruskal-Wallis test revealed differences between groups (H= 9.761) **p<0.01. **c)** Proportion of time spent freezing over time, by pairs of demonstrators and observers during the social interaction. Shaded area corresponds to time after tone presentation. Top-left panel corresponds to pairs of animals with observers that received SS (n= 11). Top-right corresponds to pairs of animals with observers that experienced freezing triggered by looming stimuli (n=14). Bottom-left panel corresponds to pairs of animals with observers that experienced freezing in response to 2MT exposure (n=9). Mean ± S.E.M. **d)** Freezing values per pair, during social interaction, normalized by the freezing of the demonstrator. Line denotes median values. **p<0.01; ****p<0.000.1.

This experiment reveals that the experience of freezing on its own is not sufficient to drive observational freezing. Together with the results of the previous experiments this finding strongly suggests that some form of conditioning, where freezing is paired with a painful event, must occur to transform this behavioral output into an alarm cue.

### Optogenetic triggering of freezing

Exposure to unsignaled spaced shocks constitutes the classical contextual conditioning paradigm. In this paradigm freezing is both a response and a putative conditioned stimulus that, like the contextual cues, can become associated with the shock. To rigorously test whether freezing can become a learned alarm cue i.e. the conditioned stimulus to which observers respond during the social interaction, we asked whether explicitly pairing freezing with shock could drive observational freezing. To this end we used artificial induction of freezing, such that the time of onset and duration of freezing was fixed across rats. We induced freezing for a period of 40 seconds, at the end of which shock was delivered. Importantly, we prevented rats from experiencing shock-elicited freezing by removing them from the chamber immediately after shock delivery, as in the DS shock condition (see table in Figure 1A). To trigger freezing artificially, we activated optogenetically the ventral lateral PAG (vlPAG) using channelrhodopsin, (Figure 4A and Figure S4), as activation of this area has been shown to elicit freezing [45,46] without inducing threat learning [46]. Consistent with prior reports, stimulation alone induced robust freezing but did not support any contextual fear learning (Figure 4D). We then tested rats that either experienced only optogenetically induced freezing (Stimulation), or experienced this form of freezing paired with footshock (Stimulation + Shock). We found that a single pairing of optogenetically induced freezing with shock was sufficient to elicit observational freezing during the social interaction, but optogenetic stimulation alone was not (Figure 4B). Indeed normalized freezing scores for the Stimulation + Shock group were different from one (p= 0.0078), showing that freezing by the observer constituted a significant fraction of freezing displayed by the dyad, whereas for stimulation only that was not the case (p>0.9999). Comparing normalized freezing across conditions with a Mann-Whitney U test revealed a significant difference between the two (U= 4, p= 0.0015) (Figure 4C).

**Figure 4.**
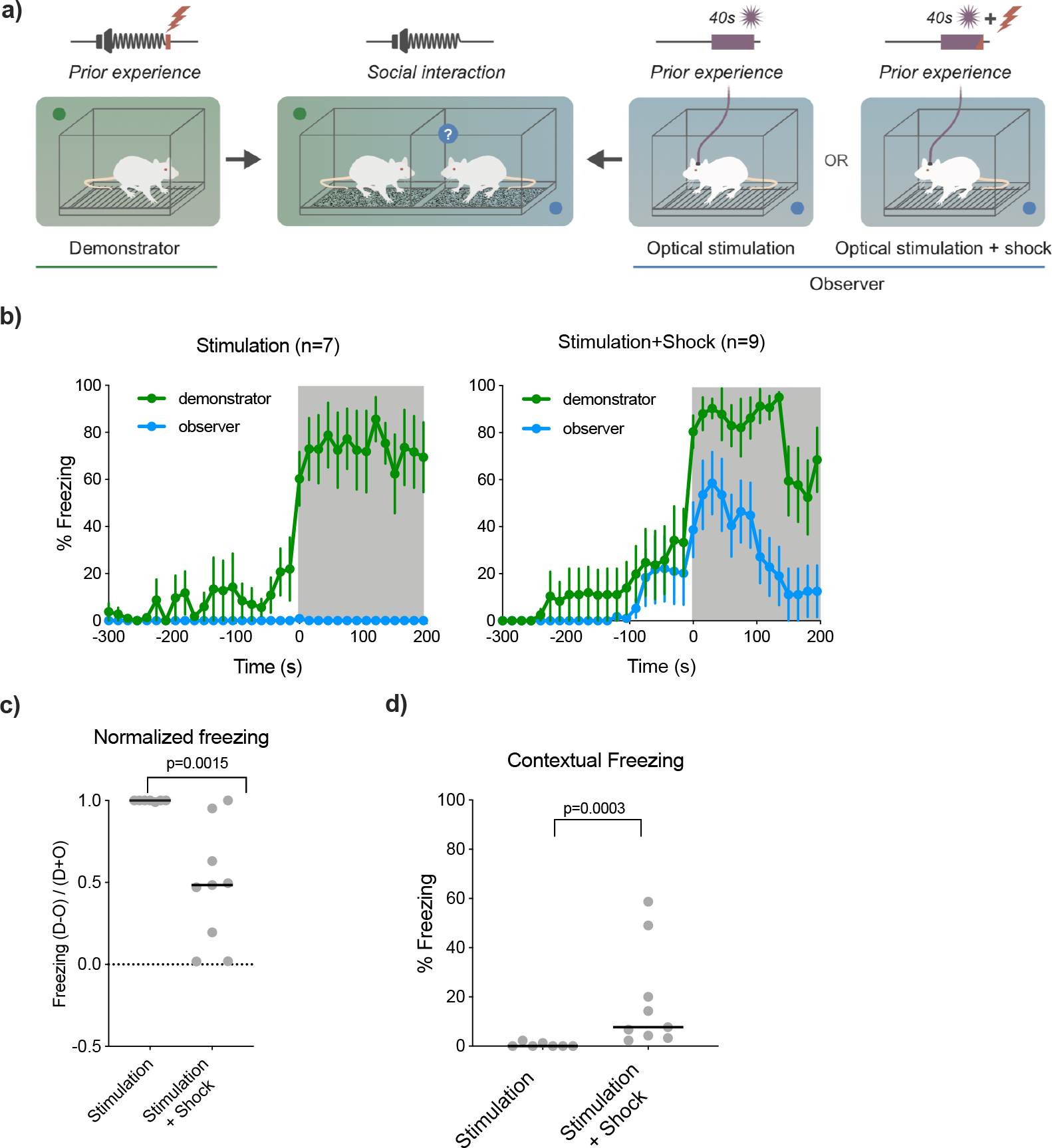
*Direct association between freezing and shock leads to observational freezing*. **a)** Schematic of the behavioral protocol using optogenetic stimulation to drive freezing. **b)** Proportion of time spent freezing over time, by pairs of demonstrators and observers during the social interaction. Shaded area corresponds to time after tone presentation. Left panel corresponds to pairs of animals with observers that received stimulation paired with shock (n= 9). Right panel corresponds to pairs of animals with observers that experienced stimulation only (n=7). Mean ± S.E.M. **c)** Freezing values per pair, during social interaction, normalized by the freezing of the demonstrator. Line denotes median values. **p<0.01. **d)** Time spent freezing in the context where animals received optogenetic stimulation, 24 hours after social interaction, as a measure of threat learning. ***p<0.001.

These data support the hypothesis that freezing becomes a conditioned stimulus to which observer animals respond during the social interaction.

## DISCUSSION

Observational freezing, triggered upon the display of the same response by a conspecific, has been shown to depend on prior experience with shock. In this study we dissected the components of prior experience that contribute to observational freezing. When receiving shocks rats experience pain, stress, their own behavioral responses and the environment in which shocks were delivered, all of which could contribute to the ability of rats in using freezing by others as an alarm cue. When we tested the role of stress or pain, triggered by shock delivery, we found that, by themselves, these do not lead to observational freezing. In addition, our results show that threat learning, whereby rats learn that the context in which they where shocked is dangerous, does not convert freezing by others into an alarm cue. Experiencing freezing triggered by non-painful stimuli that can drive innate freezing, also failed to allow the use of this behavior, when displayed by others, as an alarm cue. Finally, we show that auto-conditioning, in the form of a learned freezing-shock association, mediates observational freezing.

When investigating the contribution of prior experience with freezing to observational freezing, we found that exposure to looming shadows or exposure to 2MT [41–44], that drive innate freezing, failed to induce robust threat learning, as measured by conditioned freezing to the context. It has been shown that some predator odors, such as the odor of cat fur, are able to support contextual fear conditioning, but others, like TMT (from which 2MT is derived), are not [40]. Nonetheless, TMT exposure was shown to produce both unconditioned [47] and conditioned avoidance [48]. To our knowledge no other studies tested the ability of visual looming stimuli to reinforce threat learning, but further investigation is necessary to clarify this issue. Still, dissociation between the ability to drive strong freezing and learning has been shown using artificial stimulation of various brain regions in a study by Kim and colleagues [46]. Stimulating the basolateral amygdala, the ventrolateral or dorsal peri-acqueductal gray results in freezing, but only the later (dPAG) is able to reinforce threat learning. Together, the current experiments and this study [46] demonstrate an interesting dissociation between the ability to induce strong defensive behaviors and the ability to drive threat learning. Importantly, this supports our finding that in threat learning does not occur without shock, without which freezing does not become an alarm cue.

The main finding in this study, that rats learn to associate their own freezing with shock such that later they can use the freezing of others as a conditioned cue, raises the question of what is being associated with shock, when we induce freezing. Previously we have shown that rats use an auditory cue, the cessation of movement-evoked sound, to detect freezing by others [24]. Hence one possibility is that when rats freeze, upon shock delivery, they detected the cessation of the sound that was being produced by their own movement and associate it with the succeeding shock. In essence this would constitute a form of auditory threat conditioning. Alternatively, rats may have a representation of freezing, either a proprioceptive representation of freezing, or a command for freezing in the form of an efferent copy, which could become associated with shock. This later scenario requires rats to know that immobility is always accompanied by cessation of movement-evoked sound, which they can learn throughout their lives, such that they can use this cue to detect freezing in themselves or others. In our experimental conditions observational freezing took place either after multiple shocks, such that the first elicited freezing and the next reinforced learning, or with artificially-induced freezing followed by shock. In the wild, it is very likely that animals freeze before being attacked by the predators, such that freezing can precede the reinforcing stimulus. Therefore, the temporal relationship between freezing and shock in our experiments can reproduce a situation occurring in the wild.

In summary, we have shown that auto-conditioning, which in our paradigm occurs through the freezing-shock association, can mediate observational freezing, i.e. the ability of rats to use freezing by others as an alarm cue. Hence, a single encounter with a threat may allow rats to use information from the behavior of others to avoid danger, without the need to learn from self-experience the specific cues that predict each different form of threat. This work provides experimental evidence for a long proposed important process involved in the ability of animals to use social information.

This study shows how a single learning experience expands the repertoire of natural cues that animals can use to detect threats, while opening a new path to study how learning by self-experience comes to modulate social interactions.

## ACKNOWLEDGEMENTS

We thank the vivarium, histology and imaging services of the Champalimaud Research; Gil Costa for the illustrations in Figure 1A and Figure 2A, 3A and 4A; Ana Pereira, Susana Lima and Moita lab members for helpful discussions and comments on the manuscript. Funding: This work was supported by Fundação Champalimaud, ERCStG337747-CoCO. A.C. was further supported by FCT SFRH / BD / 51261 / 2010.

## AUTHOR CONTRIBUTIONS

A.C and M.A.M. designed all experiments. A.C., M.H. performed the experiments and analyzed the data. C.M. designed and performed, with A.C. the corticosterone measurements. A.C. and M.A.M. wrote the paper. All authors contributed in editing the final manuscript.

## STAR METHODS

### KEY RESOURCES TABLE

| REAGENT or RESOURCE | SOURCE | IDENTIFIER |
| --- | --- | --- |
| **Antibodies** |  |  |
| antibody raised in rabbits against corticosterone–carboximethyloxime-BSA | Dr. G. Makara<br>(Institute of Experimental Medicine, Budapest, Hungary) | N.A. |
| **Bacterial and Virus Strains** |  |  |
| AAV2/9. EF1a.GhR2-YFP.WPRE.hGH (Stanford), 1.58 GC/ml | University of Pennsylvania (UPENN) | AV-9-PV1522 |
| **Chemicals, Peptides, and Recombinant Proteins** |  |  |
| 2MT | Sigma-Aldrich | 2346-00-1 |
| Isoflurane | Virbac | QN01AB06 |
| Dolorex: Butorphanol Tartrate | Intervet | 42408-82-2 |
| MediGel | Clear H2O | 74-05-5022 |
| 125 I-carboximethyloxime- tyrosine-methylester | ICN-Biolink 2000 | 850489 |
| Synthetic corticosterone | Sigma | C2505 |
| **Experimental Models: Organisms/Strains** |  |  |
| Sprague Dawley Rats | Harlem | OFA-SD |
| Sprague Dawley Rats | Charles Rivers | OFA-SD |
| **Software and Algorithms** |  |  |
| Freezescan | Clever Sys | Version 2.0 |
| Bonsai Framework | Developed in-house | Version 2.3<br>https://bonsai-rx.org/ |
| PsychoPy | Peirce, 2008 | N/A |
| PRISM 8 | GraphPad | https://www.graphpad.com |
| **Other** |  |  |
| Forced Swim Cylinder | Gravoplot | N/A |
| Conditioning/optogenetic chambers | Coulbourn Instruments | H10-11R-TC |
| Shock floor of metal bars | Coulbourn Instruments | H10-11R-TC-SF |
| Sound isolation chamber | Action, automation and controls, Inc. | N/A |
| Precision programmable shocker | Coulbourn Instruments | H13-16 |
| Sound generator | Tucker- Davies Technologies | RM1 |
| Horn tweeter | VISATON | TL16H80HM |
| Microphone | Brüel and Kjaer | 4189 |
| Sound analyzer | Brüel and Kjaer | 2250 |
| Social Interaction/2MT exposure boxes | Gravoplot | N/A |
| Social Interaction/optogenetic sound attenuation boxes | Built in-house | N/A |
| Looming exposure box | Built in-house | N/A |
| LED projector | Optoma Europe Ltd, United Kingdom | ML750e |
| PointGrey/FLIR camera | Integrated Imaging Solutions GmbH, Germany | N/A |
| 200mW, 473nm laser | Changchun New Industries Optoelectronics Tech. Co., LTD | N/A |
| Stereotaxic frame | David Kopf Instruments | N/A |
| Guide Cannula | Plastics One | C315 |
| Infusion Cannula | Plastics One | C316 |
| 1µl microsyringe | Hamilton | 86200 |
| Infusion Pump | Harvard Apparatus | PHD200 |
| Optical fibers | Doric Lenses | MFC_200/245-0.37_8.5mm_RM3_FLT |
| Optical cables | Doric Lenses | MFP 105/125/LWMJ-0.22 FC-FC |
| EDTA capillary tubes | Sarsted, Granollers, Spain | N/A |
| Stereoscope | Zeiss | Zeiss SteREO Lumar.V12 |

### CONTACT FOR REAGENT AND RESOURCE SHARING

Further information and requests for resources and reagents should be directed to and will be fulfilled by the Corresponding Author, Marta Moita.

### EXPERIMENTAL MODEL AND SUBJECT DETAILS

#### Rats

Naïve male Sprague Dawley rats were obtained from a commercial supplier (Harlan, for experiments performed at the Instituto Gulbenkian de Ciência and Charles River, France, for experiments performed at the Champalimaud Foundation). After arrival animals were pair-housed in single Plexiglas top filtered ventilated cages and maintained on a 12h light/dark cycle with lights off at 8:00 p.m. All animals had *ad libitum* access to food and water.

For experiments with no surgical procedure animals weighted 250-300g on arrival. For experiments with surgical procedure (optogenetic manipulation) animals weighed 200-250g on arrival.

After surgical procedure rats were transferred with their cage-mates to double sized top-filtered cages with a buddy barrier (a perforated Plexiglas barrier covered with 1cm holes spaced at 1,5cm).

Before experimental manipulation, rats were acclimated for a minimum of one week and were handled for a variable number of sessions, until they were comfortable being held by the experimenter. All behavioral procedures were performed during the light phase of the cycle.

The Instituto Gulbenkian Ciência and the Champalimaud Foundation follow the European Guidelines for animal care. The use of vertebrate animals in research in Portugal complies with the European Directive 86/609/EEC of the European Council.

### METHOD DETAILS

#### Behavioral Apparatus

<u>Forced swim:</u> the forced swim apparatus consisted of a Plexiglas cylinder tank with 20cm diameter and 40cm high (Gravoplot). The tank was filled with 24°C water, measured with a hand held glass mercury thermometer, and kept at a level of 25 cm, to ensure that the rats could not escape the tank but also could not touch the floor while maintaining the head outside of the water.

<u>Conditioning and neutral boxes:</u> two distinct chambers (A and B), located in the same procedure room were used in a counterbalanced manner (i.e.- animals that were conditioned in chamber A were exposed to chamber B as a neutral box and vice-versa). The conditioning chambers (model H10-11R-TC, Coulbourn Instruments) were equipped with a shock floor of metal bars (model H10-11R-TC-SF, Coulbourn Instruments). The sidewalls of chamber A were made of clear Plexiglas and were cleaned with rose scented detergent. The sidewalls of chamber B were made of polished sheet metal and were cleaned with a natural soap scented detergent. The boxes were placed inside a sound isolation chamber (Action, automation and controls, Inc.) with either white (chamber A) or black (chamber B) walls. A precision programmable shocker (model H13-16, Coulbourn Instruments) delivered foot-shocks. The tones for the conditioning of the demonstrators were produced by a sound generator (RM1, Tucker-Davis Technologies) and delivered through a horn tweeter (model TL16H80HM, VISATON). The sound was calibrated using a Brüel and Kjaer microphone (type 4189) and a sound analyzer (hand held analyzer type 2250). To reduce the levels of generalization the animals were exposed to neutral boxes. For this purpose the conditioning chambers were used as neutral boxes, in a counterbalanced manner, so animals conditioned in chamber A were exposed to neutral chamber B, with the following modifications: the rod floor was covered with an acrylic plate, and the house light was on. The rats’ behavior was tracked by a video camera mounted on the ceiling of each attenuating cubicle. A surveillance video acquisition system was used to record and store all videos on hard disk and freezing behavior was posteriorly scored manually.

<u>Social interaction boxes:</u> the boxes consisted of a two partition chamber made of clear Plexiglass walls (60cm wide × 34cm height × 27cm depth) (Gravoplot), and were divided in two equal halves by a clear Plexiglas wall with 0.7cm wide vertical slits separated by 1.5cm, that allowed the animals to see, hear, smell and touch each other. Each side of the box floor contained a removable tray with bedding (the same used in the animal’s home cages). These boxes and trays were cleaned with water and ethanol 70%. The social interaction boxes were placed inside sound attenuation chambers (80cm wide × 52.5cm height × 56.5cm depth) made of MDF lined with high-density sound attenuation foam (MGO Borrachas Técnicas) and a layer of rubber. Two house lights were used inside the sound-attenuating chamber. The behavior of the animals was tracked by video cameras mounted on the walls of the sound attenuating chambers, one on each partition. A surveillance video acquisition system was used to record and store all videos on hard disk and freezing behavior was automatically scored using FreezeScan V2.0 from Clever Sys.

<u>Looming exposure box:</u> the behavioral box was made of black Plexiglas floor with dark red sides (30cm wide × 50cm height × 55cm depth) and was cleaned with a 70% ethanol solution. This box was placed in a room with ceiling lights on. Stimuli were projected with an LED projector (ML750e, Optoma Europe ltd, United Kingdom) onto an opaque white Plexiglas screen placed on top of the behavioral box. The behavior was captured with an infrared camera PointGrey/now FLIR Integrated Imaging Solutions GmbH, Germany) controlled by a custom workflow using the Bonsai visual programming language [49] and stored on hard disk for posterior manual scoring.

<u>2MT exposure box:</u> this box was made of clear Plexiglas walls (60cm wide × 34cm height × 27cm depth) (Gravoplot), and was cleaned with a 70% ethanol solution. Because the nature of the stimulus used, the 2MT experiments were performed inside a fume hood. The behavior was recorded with a handheld video camera, and the videos stored on hard disk for posterior manual scoring of freezing.

<u>Optogenetic box:</u> one conditioning chamber (model H10-11R-TC, Coulbourn Instruments), equipped with a shock floor of metal bars (model H10-11R-TC-SF, Coulbourn Instruments) was placed inside a custom made sound attenuating chamber 90cm wide × 45cm height × 52.5cm depth) made of MDF lined with high-density sound attenuating foam (MGO Borrachas Técnicas). A precision programmable shocker (model H13-16, Coulbourn Instruments) was used to deliver foot-shocks. A surveillance video acquisition system was used to record the conditioning session and the videos were stored on hard disk. The blue light was deliverer by a 200mW, 473nm laser (Changchun New Industries Optoelectronics Tech. Co., LTD).

#### Surgery

<u>Virus injection and fiber implant:</u> Animals were anaesthetized with Isoflurane (4% for induction, 2% for maintenance Vetflurane 1000mg/m, Virbac) and placed in a stereotaxic frame (David Kopf Instruments). Small craniotomies were made using standard aseptic techniques. Animals were targeted bilaterally to the PAG (stereotaxic coordinates from Bregma, anterior-posterior: −7.8mm, dorsal-ventral: −6; medial-lateral: 2.9 with the stereotaxic arm angled at 20.14°; Paxinos and Watson 2007) using stainless steel guide cannulas (24 gauge; Plastics One). Following guide cannula placement, 0.3 μl injections of the virus were made through a stainless steel injection cannula (31 gauge: Plastics One), which protruded 2.0 mm beyond the tip of the guide cannula and was attached to a Hamilton syringe via polyethylene tubing. Injections were made at a rate of 0.02 μl/min, controlled by an automatic pump (PHD 2000; Harvard Apparatus) and the injector was left in place for 10 min post-injection to ensure correct dispersion of the virus. After injection, cannulas were removed and optical fibers (200μm, 0.37NA, Doric lenses) were implanted targeting the same coordinates as virus injections, and affixed to the skull using stainless steel mounting screws (Plastics One) and dental cement (TAB 2000, Kerr). Animals were kept on a heating pad throughout the surgical procedure. Post-operative care included subcutaneous injection of 0.3 mL of Dolorex (Butorphanol Tartrate, 2mg/kg, Intervet) for post-operative immediate analgesia purposes and a supplement of food with Carprofen 5mg/kg/day (MediGel CFP, Clear H2O) for pain management of post-operative day. Rats were kept for 3 weeks before behavioral manipulation to allow for maximal expression of virus.

#### Viral vectors

Adeno-associated virus containing Chr2 (AAV2/9. EF1a.ChR2-YFP.WPRE.hGH (Stanford), 1.58 GC/ml was produced by and purchased from University of Pennsylvania (UPENN) vector core facility.

#### Behavioral procedures

Experiments with social interactions were done with pairs of cage-mate rats, where one of them was randomly assigned to be the demonstrator and the other the observer. On the first two experimental days each rat was exposed for 15 minutes to each of the boxes, conditioning, social and neutral, with time between exposures ranging from 5 to 24 hours. Animals that were in the forced swim group did not get exposed to the conditioning box. In the experiment for the investigation of the corticosterone changes in response to prior experience, the animals were also pair-housed. No demonstrators or observers were assigned in this procedure, and both cage-mates underwent the same experience (i.e. each member of the pair had the same experience either forced swim or shocks).

<u>Manipulation of stress experience:</u> On the third day, after pre-exposures, demonstrator (DEM) rats were placed in the conditioning chamber. After an initial period of 5 minutes, demonstrator rats received 5 tone-shock pairings (tone: 15s, 5kHz, 70dB; shock 1mA, 1s), with the tone and shock co-terminating and an average inter trial interval of 180 seconds (ranging from 170 to 190s). After the last tone-sock pairing animals were returned to their home cage. Observer animals were placed either in the conditioning chamber or the forced swim apparatus, and were subjected to the protocols that correspond to the different condition of their prior experience. Spaced Shock observers (OBS-SS) received 3 unsignaled shocks (with the same shock intensity and schedule as demonstrators) after which they returned to the home cage. Forced Swim observers (OBS-FS) were subjected to 15 minutes immersion in water at 24°C. After this period rats were helped out of the water with a wire mesh and scrubbed with a towel until dry and warm, and returned to their home cage.

On the fourth day, the different pairs of rats, demonstrators and observers from the spaced shock protocol group (DEM-SS/OBS-SS), and demonstrators and observers from the forced swim protocol group (DEM-FS/OBS-FS), were tested in the social interaction box. Each animal was placed on one side of the two-partition box, and after a 5 minutes baseline period 3 tones (same tone as described above) were presented, with a 3 minutes inter-trial interval. The behavior of both animals was recorded for offline scoring of freezing.

On the fifth day, observers that were conditioned (OBS-SS) were placed back in the conditioning chamber and their behavior was recorded for posterior assess of context fear by scoring the time they spend freezing over a period of 5 minutes.

<u>Corticosterone measurements:</u> Animals underwent behavior protocols similar to the ones described above for the OBS-SS and OBS-FS, except there was no exposure to the social interaction box since no social interaction was performed. One day after exposures blood samples were taken by tail-nick, to obtain basal levels of circulating hormone and to habituate the animals to the sampling procedure. Basal corticosterone levels were measured 24hours before the behavioral procedures, in order to avoid interference with the measures taken after exposure to the different stressors. On the prior experience day blood samples were collected by tail-nick for both groups with the following schedule: immediately after the exposure to the behavioral protocol (‘stressor’ time point) to investigate the acute stress response of each treatment, and at two time points after- 30 minutes (post 30) and after 60 minutes (post 60), to establish the recovery profiles. The tail nick consisted of gently wrapping the animals with a cloth, making a 2 mm incision at the end of one of the tail artery and then massaging the tail while collecting 300 μl of blood, within 2 minutes, into ice-cold EDTA capillary tubes (Sarsted, Granollers). Plasma obtained after centrifugation was stored at −30 °C until assay. Plasma corticosterone levels were determined by double-antibody radioimmunoassay (RIA) procedures. Corticosterone RIA used ^125^ I-carboximethyloxime- tyrosine-methylester (ICN-Biolink 2000, Spain) as the tracer, synthetic corticosterone (Sigma) as the standard and an antibody raised in rabbit against corticosterone-carboxi- methyloxime-BSA kindly provided by Dr. G. Makara (Institute of Experimental Medicine, Budapest, Hungary). The RIA protocol followed was recommended by Dr. G. Makara (plasma corticosteroid-binding globulin was inactivated by low pH) (Bagdy and Makara, 1994). All samples to be compared were run in the same assay to avoid inter-assay variability.

<u>Experience with shock:</u> These experiments were performed in the facilities of Instituto Gulbenkian de Ciência (all other experiments were performed at the Champalimaud Center for the Unknown). Experiments were performed with pairs of cage-mate rats, each one assigned randomly to be the demonstrator or the observer. On the first two experimental days each rat was exposed for 15 minutes to each of the boxes, conditioning, social and neutral, with time between exposures ranging from 5 to 24 hours, except for Immediate Shock (IS) observers that are exposed only to the social interaction and neutral boxes.

On the third day, demonstrator (DEM) rats were placed in the conditioning chamber. After an initial period of 5 minutes, demonstrator rats received 5 tone shock pairings (tone: 15s, 5kHz, 70dB; shock 1mA, 1s), with the tone and shock co-terminating and an average inter trial interval of 180 seconds (ranging from 170 to 190s). After the last tone-sock pairing animals were returned to their home cage. Observer animals were placed in the conditioning chamber and were subjected to the protocols that correspond to the different condition of their prior experience. Spaced Shock observers (OBS-SS) received 3 un-signaled shocks (with the same shock intensity and schedule as demonstrators) after which were returned to the home cage. Immediate Shock observers (OBS-IS) were taken to the conditioning room alone, in a small cage through a different pathway from the holding room, to avoid generalization or conditioning to any part of the procedure. Once they entered the conditioning chamber they received immediately 3 un-signaled shocks (1mA, 0.5s) with 100ms interval after which they returned to their home cage. Delayed Shock observers (OBS-DS), after a 5 minutes baseline, receive 3 un-signaled shocks (1mA, 0.5s) with 100ms interval after which they return immediately to their home cage.

On the fourth day, the different pairs of rats were tested in the social interaction box. Each animal was placed on one side of the two-partition box, and after a 5 minutes baseline period 3 tones (same tone as described above) were presented, with a 3 minutes inter-trial interval. The behavior of both animals was recorded for offline scoring of freezing.

On the fifth day, observers were placed back in the conditioning chamber and their behavior recorded for posterior assess of context fear by scoring the time they spend freezing over a period of 5 minutes.

<u>Experience with freezing:</u> these experiments were performed at the facilities of the Champalimaud Foundation, and followed the same behavior procedures and schedule described above, in exposures, conditioning, social interaction and context test, except for the conditioning day where looming and 2MT observers received a different prior experience: observers of the Looming (Loom) group were placed in the box for a period of 5 minutes to establish baseline. The looming stimulus was generated with PsychoPy (Peirce, 2008), consisting of an expanding black dot (0cm to 30cm diameter in 0.5 seconds), on a grey background. The stimuli were presented every second, with 20 stimuli per looming session and a 30 second interval between looming sessions. A total of 8 looming sessions was presented in 380 seconds. Animals were then returned to their home-cage. Observers subjected to the 2MT presentation were taken in single boxes to the room with the fume hood. After a 5 minutes baseline, 3 small filter papers embedded with 6 μl of 2MT were presented to the animal, with an inter-stimulus interval of 2 minutes, inside a small plastic container. 2 minutes after the last presentation the animal was returned to the transport box and then transferred to his home-cage.

<u>Optogenetic freezing:</u> experiments were performed with pairs of cage-mate rats, each one assigned randomly to be the demonstrator or the observer. On the first two experimental days each rat was exposed for 15 minutes to each of the boxes, conditioning, social and neutral, with time between exposures ranging from 5 to 24 hours.

On the third day, demonstrator (DEM) rats were placed in the conditioning chamber. After an initial period of 5 minutes, demonstrator rats received 5 tone shock pairings (tone: 15s, 5kHz, 70dB; shock 1mA, 0.5s), with the tone and shock co-terminating and an average inter trial interval of 180 seconds (ranging from 170 to 190s). After the last tone-sock pairing animals were returned to their home cage. For observers, two fiber optic cables (200μm, 0.22 NA, Doric lenses) terminating in ferrules were attached to the chronically implanted optic fibers. The rats were then placed in the conditioning chamber. After a baseline period of 3 minutes, 40 seconds of light stimulation were delivered (20mW, 20Hz, 40ms pulses). The animals froze for the entire duration of the stimulation that co-terminated with one shock (1,5mV, 1,5 seconds) only for the Stimulation + shock group. After the session the optic cords were removed and the animals returned to their home cage.

On the fourth day the pairs of were tested in the social interaction box. Each animal was placed on one side of the two-partition box, and after a 5 minutes baseline period 3 tones (same tone as described above described) were presented, with a 3 minutes inter-trial interval.

On the fifth day, observers were placed back in the conditioning chamber and their behavior recorded for posterior assess of context fear by scoring the time they spend freezing over a period of 5 minutes.

#### Histological processing

Animals were deeply anesthetized with pentobarbital (600 mg/kg, i.p.) and transcardially perfused with PBS (0.01M), followed by ice-cold 4% paraformaldehyde in 0.1M phosphate buffer (PFA). Brains were removed and post-fixed overnight in a 4% PFA solution at 4°C. The brains were then transferred to a 30% sucrose solution in PBS (0.01M) and kept at 4°C until sunken. Coronal sections of 50 μm containing PAG (for viral expression and fiber placement verification) were cut, collected on coated slides and mounted using mowiol. A stereoscope (Zeiss Lumar) was used to examine the slides.

#### Statistical analysis and exclusions

Statistical analyses were performed with the software PRISM 8 (GraphPad).

<u>Corticosterone investigation experiment:</u> the normality of the data was evaluated by means of the Kolmogorov-Smirnov test. To compare the basal levels of the hormone, a Mann Whitney U test was performed. To analyze the corticosterone changes in relation to the protocol experienced by the animal, a repeated measures 2way-ANOVA was performed, with treatment (forced swim or spaced shocks) as between-subject’s factor and time (basal levels, after experience levels and 30 and 60 minute post experience levels) as the within-subject’s factor. Post-hoc analysis was done using the Holm-Sidak’s test for multiple comparisons. Significance level was set at p<0.05.

<u>Criteria for inclusion of data for analysis:</u> since we are investigating how previous experience influences a behavior that is triggered by the behavior of another animal, we decided to exclude from our analysis pair of animas where the demonstrator failed to display defensive responses. To this end, we divided the period after the presentation of the tone (tone that the demonstrators were conditioned to fear and thus that should trigger defensive responses such as freezing) in two 90-second parts. Pairs were excluded if demonstrator animals would not display freezing for at least 33% of this time on either parts of the post tone presentation.

For the stress manipulation experiment a total of 29 pairs of animals was used- 13 with SS observers and 16 with FS observers. From these 3 and 4 pairs were excluded from the respective groups. Final n: SS= 10; FS= 12.

For the shock manipulation experiment a total of 44 pairs of animals was used- 12 with SS observers, 17 with IS observers and 15 with DS observers. From these 1, and 2 pairs were excluded from the SS and IS groups, respectively. Final n: SS= 11; IS= 15; DS=15.

For the freezing manipulation experiment a total of 49 pairs of animals was used- 14 with SS observers, 20 with “Looming” observers and 15 with "2MT" observers. From these 3, 6 and 6 pairs were excluded from the respective groups. Final n: SS= 11; Loom= 14; 2MT= 9.

For the optogenetic manipulation experiment a total of 24 pairs of animals was used- 11 with “Stimulation+shock” observers, and 13 with “Stimulation” observers. From these 2 and 5 pairs were excluded from the groups, respectively. From the “Stimulation” group, an additional pair was excluded because the observer was freezing before stimulation during training. Final n: Stimulation+Shock= 9; Stimulation= 7.

<u>Experience with shock:</u> to test the robustness of the paradigms we use for prior experience of observers we performed a context test on all of the observers. We scored the percentage of time they spent freezing in the context where they had received the un-signaled shocks. Since the distribution of these values did not comply with the normality assumption we investigated the differences between groups using the non-parametric Kruskal-Wallis analysis with post-hoc Dunn’s test for multiple comparisons. Significance was set at p<0.05.

<u>Experience with freezing:</u> to test the sufficiency of experiencing freezing as a condition for observational freezing we exposed animals to visual looming stimuli and to the predator odor 2MT. We needed to make sure that our observers were in fact freezing during prior experience. For that we scored the time each animal spends freezing during training i.e. prior experience. We investigated the differences between groups via a Kruskal-Wallis test with post-hoc Dunn’s test for multiple comparisons. Significance was set at p<0.05.

<u>Optogenetic stimulation:</u> all animals that reliably froze during the entire stimulation were included in the analyses. Still, we performed histological verification of fiber optic placement and viral expression. All animals had viral expression; most animals had both fibers in the vlPAG, some at the border with the dorsal raphe. Two animals had unilateral optic fiber placement. To test if the optogenetic stimulation alone was not aversive per se, and capable of inducing threat-learning, we performed a context test and measured the time spent freezing in the stimulation box. Since the distribution of these values did not comply with the normality assumption we investigated the differences between groups using the non-parametric Mann-Whitney U test. Significance was set at p<0.05.

<u>Normalization:</u> we performed a normalization of the freezing time to get one value per pair of animals in order to compare groups. The normalization was done by calculating the difference in freezing by the observer and by the demonstrator in a dyad over the total amount of freezing by the pair ((d-o)/(d+o)) the normalized values vary between −1 and 1, with values closer to zero meaning that both animals contributed equally to the freezing of the pair, and values closer to 1 or −1 reveal that the demonstrator, or the observer respectively, contributed with most of the freezing. The normalized values were then used to investigate if there is any difference between groups using a Mann-Whitney U test for the FS/SS and for the optogenetic groups comparison, and a Kruskal-Wallis analysis with post-hoc Dunn’s test for multiple comparisons for the shock and freezing manipulations. Significance was set at p<0.05.

