## Supplementary Figures for "Freezing displayed by others is a learned cue of danger resulting from co-experiencing own-freezing and shock"

**Figure S1**

**a)**

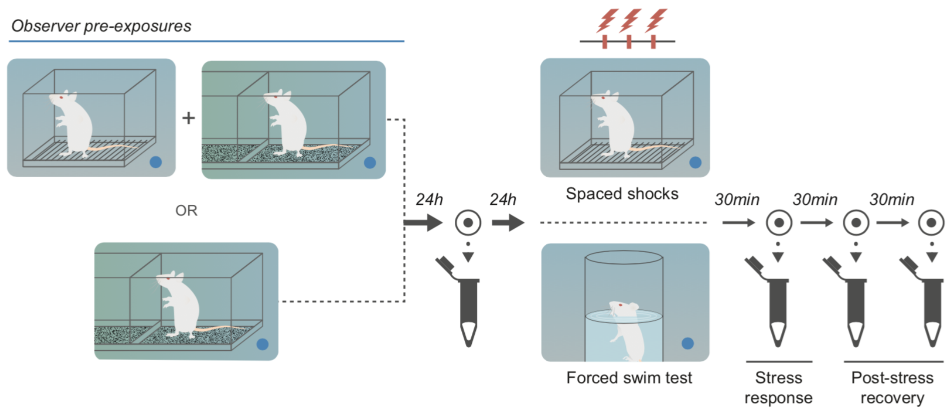


**b)**

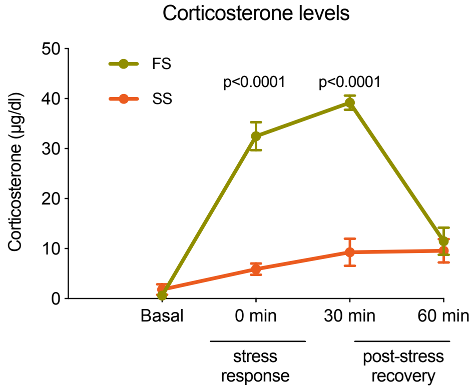


**c)**

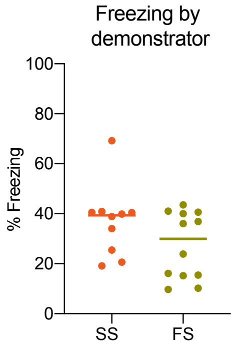


**Figure S1.** *Corticosterone levels in animals that experience SS versus FS. Related to Figure 1.*

**a)** Schematic of the timeline of blood sample collection.
**b)** Plasmatic corticosterone dynamics (μg/dl) of animals that experience Spaced Shocks (SS, n=8) and animals that experience Forced Swim (FS, n=8). Basal levels were measured 24 hours before behavioral procedures. Mann-Whitney U test revealed no difference between groups ($U= 29$, $p= 0.775$). Time zero measures the stress response immediately after the behavioral procedure. Two additional samples were taken in the post-stress recovery period at 30 and 60 min after behavioral procedure (see timeline in a)). The 2way-ANOVA revealed different corticosterone responses to the FS treatment compared to the SS protocol. Significant effects of treatment ($F(1,14)= 54.87$, $p<0.0001$), sampling time ($F(2,28)= 36.86$, $p<0.0001$) and interaction between treatment by sampling time ($F(2,28)= 44.85$, $p<0001$) were found. Subsequent analysis revealed that animals that experienced FS showed a greater corticosterone response immediately after the stress and after 30 minutes. No significant differences were found at the post 60 minutes time point. ****$p<0.0001$.
**c)** Proportion of freezing of demonstrator animals of both the SS and FS groups. A Mann-Whitney U test revealed no differences between groups ($U= 42$, $p=0.2543$).

Figure S2

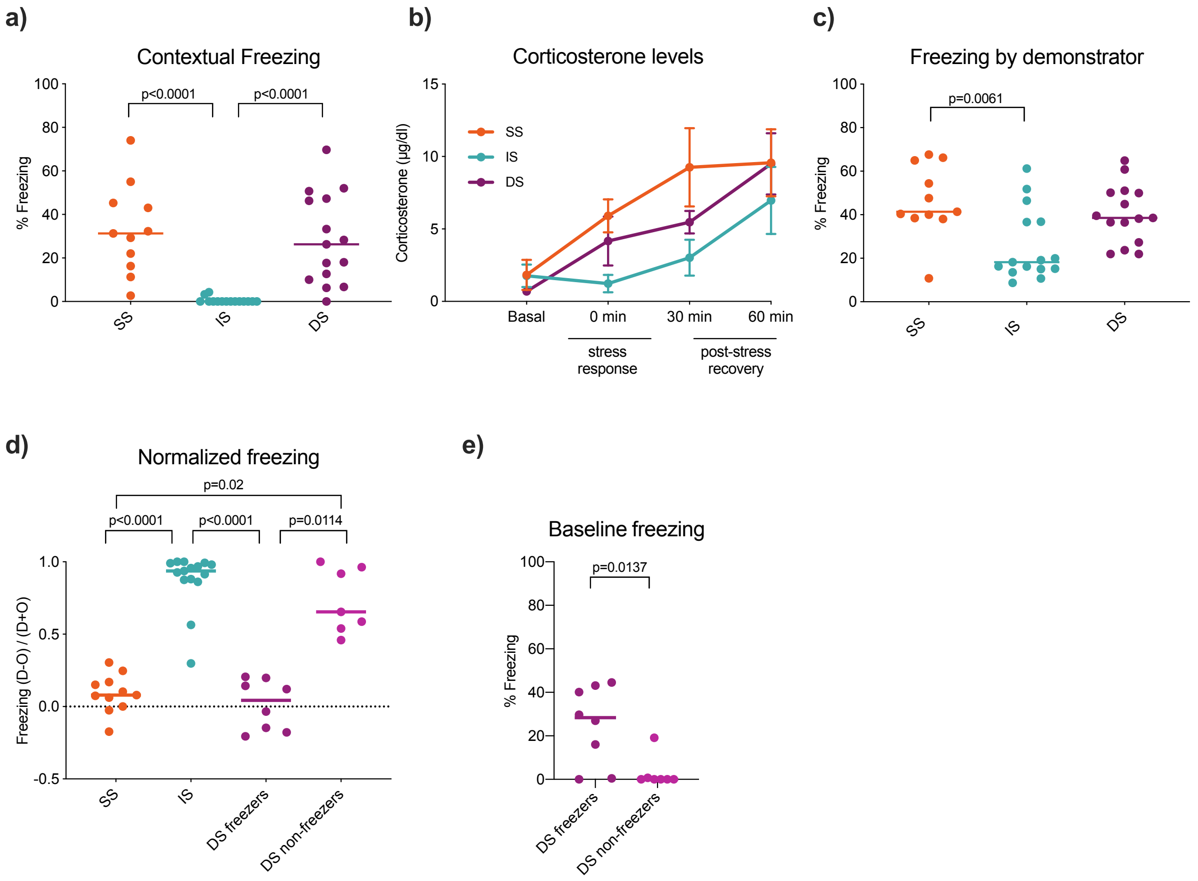


**Figure S2.** *Characterization of the different shock protocols for the prior experience of observers and normalized freezing scores after median split of the delayed shock dyads. Related to Figure 2.*

**a)** Time spent freezing in the context where animals received shocks, 24 hours after social interaction, as a measure of threat learning. A Kruskal-Wallis test revealed differences between groups (H= 26.16) ****p<0.0001.
**b)** Blood corticosterone (µg/dl) of animals that experience SS (n=8), IS (n=8) and DS (n=8) in four time points. Basal levels are measured 24 hours before behavioral procedures. Time zero corresponds to time immediately after the behavioral procedure. Times 30 min and 60 min correspond to recovery period.
**c)** Average time that the demonstrators spent freezing during the social interaction in the three experimental groups: SS (n=11), IS (n=15) and DS (n=15). A Kruskal-Wallis test revealed differences between groups (H= 10.57) **p<0.01.
**d)** Freezing values per pair, during social interaction, normalized by the freezing of the demonstrators. SS (n=11), IS (n=15), DS freezers (n=8) and DS non-freezers (n=7). Line denotes median values. Kruskal-Wallis test (H= 30.18) *p<0.05; ****p<0.0001.
**e)** Time spent freezing during baseline for observer animals after median split of the normalized values of freezing. Mann-Whitney U test shows difference between groups (U= 7.5) *p<0.05.

### Figure S3

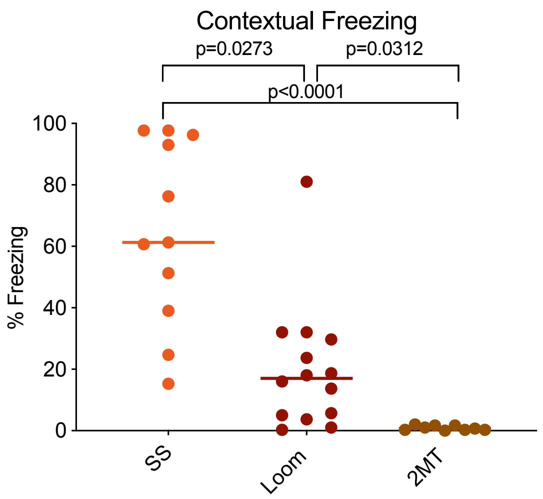


**Figure S3.** *Threat learning from observers with different freezing experiences. Related to Figure 3.*

Time spent freezing in the context where animals experienced freezing, 24 hours after social interaction, as a measure of threat learning. A Kruskal-Wallis test revealed differences between groups (H= 22.85). *p<0.05; ****p<0.0001.

**Figure S4**

a) b)

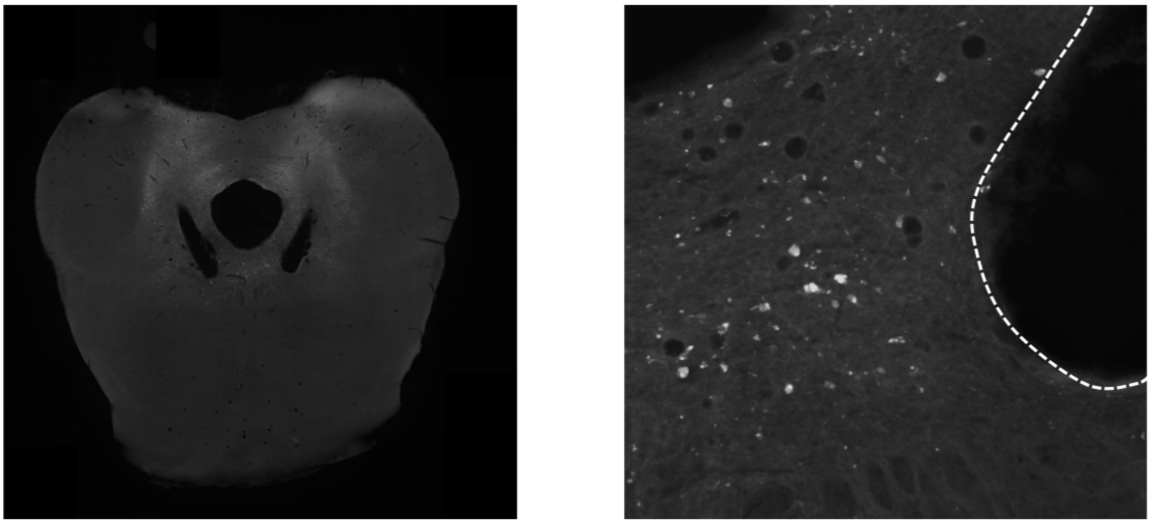

**Figure S4.** *Optical fiber placement in the vlPAG. Related to figure 4.*

**a)** Example photomicrograph showing optical fiber placement targeting the vlPAG. The inclusion of individuals was based on freezing behavior upon stimulation and not on fiber placement. These included animals with unilateral fiber placement or fiber placements close to the dorsal raphe.
**b)** Magnification showing neuronal cells infected with ChR2.
